# Adaptation-induced blindness is orientation-tuned and monocular

**DOI:** 10.1101/048918

**Authors:** Deborah Apthorp, Scott Griffiths, David Alais, John Cass

## Abstract

We examined the recently discovered phenomenon of Adaptation-Induced Blindness (AIB), in which highly visible gratings with gradual onset profiles become invisible after exposure to a rapidly flickering grating, even at very high contrasts. Using very similar stimuli to those in the original AIB experiment, we replicated the original effect across multiple contrast levels, with observers at chance in detecting the gradual onset stimuli at all contrasts. Then, using full-contrast target stimuli with either abrupt or gradual onsets, we tested both the orientation tuning and interocular transfer of AIB. If, as the original authors suggested, AIB were a high-level (perhaps parietally mediated) effect resulting from the ‘gating’ of awareness, we would not expect the effects of AIB to be tuned to the adapting orientation, and the effect should transfer interocularly. Instead, we find that AIB (which was present only for the gradual onset target stimuli) is both tightly orientation-tuned and shows absolutely no interocular transfer, suggesting a very early cortical locus.

## Introduction

Neurons at the earliest stages of visual cortical processing respond preferentially to retinal image movement (De Valois et al. 1982; Hubel and Wiesel 1962; Movshon and Newsome 1996). Not only does retinal motion inform us about the relative speeds and trajectories of objects in our visual environment (including our own bodies), it can capture our attention (Cass et al. 2011), break object camouflage, and can even inform us about the surface properties of objects (Doerschner et al. 2011).

Prolonged exposure to ‘fast’ (~10 Hz) movement or flicker, however, can temporarily alter visual processing and cause a range of perceptual effects. Movement at a recently adapted retinal location appears slowed (Thompson 1981 see Hietanen et al. 2007 for an electrophysiological analogue of this effect). Under certain conditions, adaptation can even cause illusory reversals in perceived motion direction (Arnold et al. 2014). Notably, flicker adaptation produces subsequent elevation in detection thresholds, most significantly when adaptor and target stimuli are similarly oriented (Campbell and Kulikowski 1966; Cass et al. 2012).

In 2010, Motoyoshi and Hayakawa introduced a compelling new illusion, Adaptation Induced Blindness (AIB) (Motoyoshi and Hayakawa 2010). Following prolonged exposure to ~10 Hz motion, a target grating is then ramped on from zero to full (or near full) contrast, with either a gradual or an abrupt temporal profile. The slope of this onset ramp has a profound effect on subsequent perception: whereas high contrast patterns with abrupt onsets are clearly visible (although still somewhat affected), gradually presented patterns become temporarily ‘invisible’. Motoyoshi and Hayakawa (2010) attribute this effect to relatively high-level processes, possibly involving parietal brain regions, suggesting that the visual transients associated with abrupt-onset stimuli are necessary to prompt visual awareness of the stimuli. They further suggest that even though the ‘invisible’ stimuli are not available to awareness, they can still cause low-level effects such as the tilt illusion, suggesting that there is some processing of the suppressed stimuli at lower levels of visual processing (e.g., V1). This study is designed to evaluate the relative contribution of low and high-level visual processes to the AIB phenomenon.

An image presented to the retina generates a cascade of neural activity throughout the cortex, principally via lateral geniculate nucleus (LGN) and primary visual cortex (V1). LGN neurons are monocular, receiving exclusive input from the ipsilateral eye. They are selective for parameters such as spatial and temporal frequency and retinal location, but are not orientation-selective, as their receptive fields are roughly isotropic in the orientation domain (Hubel and Wiesel 1961, 1962, 1968; Alonso et al. 2001; Cheong et al. 2013). Consequently, adapting LGN neurons to a Gabor stimulus drifting or flickering in the alpha (8 −12 Hz) range – similar to that used by Motoyoshi and Hayakawa (2010) – causes their subsequent responses to be strongly attenuated to an approximately equivalent extent across the orientation spectrum (Solomon et al. 2004).

The transmission of signals from LGN to V1 is functionally significant for several reasons. For example orientation selectivity first emerges in V1 (Hubel and Wiesel 1968; Ferster and Miller 2000). While neurons in both areas respond to luminance-defined edges, only V1 contains an abundance of highly orientation-selective cells (Ringach et al. 2002); most LGN cells respond equivalently to all orientations. As observed in LGN, profound adaptation effects are observed in V1 following prolonged exposure to flickering gratings (Boynton and Finney 2003). Consistent with their orientation tuning, and unlike in LGN, V1 adaptation effects are orientation specific (Campbell and Kulikowski 1966; Blakemore and Nachmias 1971; Anderson and Burr 1985; Snowden 1991). Therefore, if AIB is mediated at a cortical stage of processing, reports of invisibility following adaptation to rapid flicker should peak at the adapting orientation.

Another functional change between LGN and V1 is the emergence of binocularly-driven responses (Hubel and Wiesel 1962). Whereas LGN neurons are driven monocularly, V1 is the first stage in visual processing to contain neurons which respond (in varying proportions) to both eyes. Because V1 cells can be driven by either eye, many adaptation effects transfer interocularly Gilinsky:1969uu, Blake:1981ta, Bjorklund:1981vh, although some do not. For example, both Cass et al. (2012) and Baker and Meese (2012) reported that the loss of sensitivity following adaptation to a flickering Gabor of low spatial frequency (~<1.5 c.p.d.) failed to transfer when adapting and testing in different eyes (i.e. interocularly). Intriguingly, this monocular threshold elevation effect was insensitive to orientation differences between adaptor and target patterns (i.e., it was purely isotropic).

In this paper we will investigate both the orientation tuning and interocular transfer of AIB. Given that AIB and classical threshold elevation result from similar adaptation paradigms (Cass et al. 2012; Baker and Meese 2012), we might expect to observe a similar pattern of results: (i) an orientation-invariant component which is specific to the adapted eye (i.e., failing to transfer inter-ocularly), and therefore possibly mediated at precortical locus (Solomon et al. 2004); and (ii) an orientation-specific component which transfers between the eyes, and is therefore likely to be cortical in origin. By contrast, if AIB were a predominantly ‘high-level’ phenomenon, possibly involving parietal structures (Motoyoshi and Hayakawa 2010), we would expect AIB to exhibit near-complete interocular transfer.

## Experiment 1: AIB across contrast levels

Experiment 1 measured detection of target Gabors as a function of target contrast. Using the method of constant stimuli, participants adapted to fast (10Hz) flicker and in a 4AFC paradigm (a simplified version of Motoyoshi and Hayakawa’s original 8AFC paradigm), and indicated the perceived location of a gradually on-ramped target stimulus. Target stimuli were presented at contrasts of 0.06, 0.09, 0.13, 0.18, 0.25, 0.35, 0.5, 0.71 and 1 (full contrast).

### Methods

*Participants* Six adults with normal or corrected-to-normal vision participated in the experiment. All participants, including three of the authors, were experienced psychophysical observers; three were nave to the purpose of the experiment.

*Stimuli* Stimuli were programmed in MATLAB version 2015b using the Psychophysics Toolbox version 3.0.12 (Brainard 1997; Pelli 1997). Visual stimuli were presented on a ViewPixx custom LCD monitor with a screen resolution of 1920 × 1080 pixels and a vertical refresh rate of 120 Hz. The monitor was gamma-corrected to ensure linear luminance output and was controlled by a quad core Mac Pro computer. The ViewPixx monitor incorporated a digital-to-analogue converter that provided 12-bit resolution for measurement of low contrast thresholds. Maximum and minimum luminances were 96.2 and 0.1 *cd*/*m*^2^, and mean luminance was 48.05 *cd*/*m*^2^. Participants sat in a darkened room with their head supported by a chin rest at a distance of 41 cm from the monitor and made responses on a standard keyboard.

The adapting stimulus was a counterphasing grating composed of two superimposed sine-wave gratings drifting leftwards and rightwards at a speed which produced 8 Hz flicker, equal to the speed of the drifting gratings used by Motoyoshi and Hayakawa (2010), and was presented at full contrast. Counterphasing gratings were used as adaptors instead of drifting gratings because drifting gratings produce potentially confounding motion aftereffects (Pantle and Sekuler 1969), and might reasonably be expected to induce saccades in the direction of the drift, thereby disrupting fixation and adaptation. The spatial frequency and orientation of the adaptor were kept constant at 1.5 c.p.d and 0°, respectively. Each adaptor was enclosed in a circular aperture whose diameter subtended 3.92° of visual angle and was edge-blurred by a cosine ramp that transitioned from minimum to maximum over 0.78° of visual angle.

The target stimulus was a static Gabor with equal spatial frequency to the adaptors but variable contrast. The Gaussian spatial envelope of the target had a standard deviation of 0.65° of visual angle and was equal in diameter to the adaptors. The target grating ramped on and off in a temporal Gaussian window of 1000ms with a standard deviation of 200ms, starting 1000ms after adaptor offset. This was jittered by a period of 1–240ms to increase temporal uncertainty.

The visual display comprised four virtual display windows. The counterphasing adapting gratings appeared within these windows, such that the centre of each adapting grating was at 5.9° eccentricity, as in Motoyoshi and Hayakawa (2010), and appeared on a grey background held at mean luminance.

*Procedure* Participants pressed a key to initiate trials and were presented with four counterphasing adaptor gratings, one appearing in each aperture. Adaptors were displayed for 30 s for the initial two trials, with 10 s ‘top-ups’ for subsequent trials. Following offset of the adaptors, a spatial four-alternative forced-choice task required observers to use a standard keyboard to indicate which aperture the target appeared in. Before the sessions, participants were reminded to keep their eyes fixated on the central cross and to avoid blinking as much as possible. Each target contrast was presented 10 times, randomly interleaved, for two blocks, meaning each participant completed 20 trials per target contrast level, a total of 180 trials. Targets were ramped on and off in a temporal Gaussian envelope with a standard deviation of 200 ms.

The phase of the target grating was randomised and its onset was temporally jittered between 0 and 240 ms to introduce a degree of temporal uncertainty to the task. Participants were instructed to maintain fixation on the central fixation cross and to guess if they failed to observe the target. The fixation cross appeared white during presentation of the adaptors and grey during target onset and offset, before changing back to white to prompt the participant to respond. The next trial began immediately after response. Each of the adaptation blocks comprised 90 trials and took approximately 40 minutes to complete.

### Results

Although some subjects showed slightly above-chance performance (see Figure 1a), on average there was no difference in performance across contrast levels (in other words, unlike in conventional contrast adaptation, there was no increase in detection levels at higher contrasts), F(8.40) = 1.18, p = .335; see Figure 1b. In addition, there was no significant difference from chance performance (25% detection) across all the contrast levels (see Table 1).

**Figure 1.**
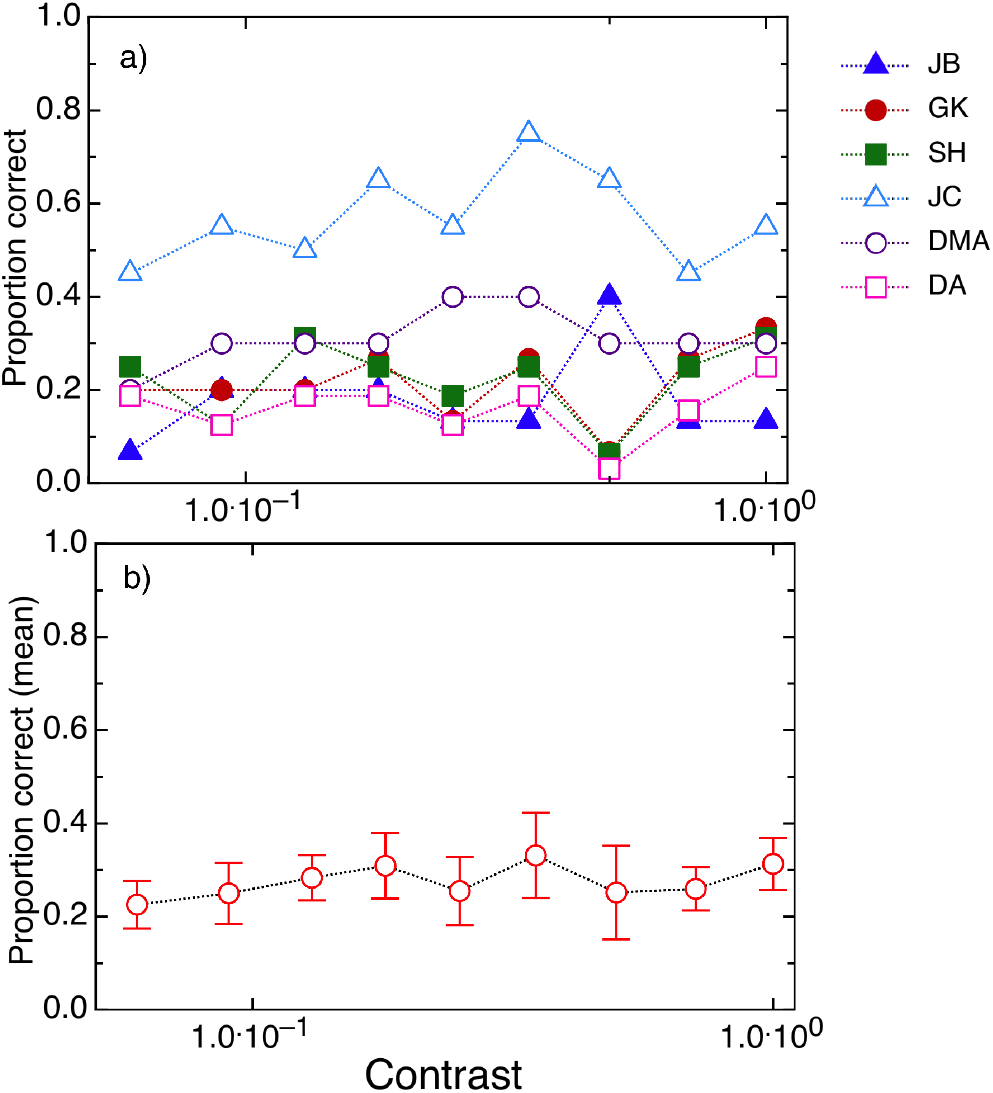
Results from Experiment 1, testing detection of gradually-onset target stimuli after adaptation to fast (8 Hz) flicker at the full range of target contrasts. (a) individual results; (b) mean results across the 6 participants (error bars show ±1 standard error). As this was a 4AFC task, chance detection threshold was .25. There was no statistical difference from chance at any of the contrast levels, suggesting full adaptation-induced blindness.

**Table 1.** t-tests comparing detection thresholds to chance (0.25) across all the contrast levels. *Note:* for all tests, the hypothesis was that the population mean was different from 0.25 (i.e., a two-tailed hypothesis). Tests are not corrected for multiple comparisons.

| Contrast level | t | df | p-value |
| --- | --- | --- | --- |
| 0.06 | -0.474 | 5 | 0.656 |
| 0.09 | 0.000 | 5 | 1.000 |
| 0.13 | 0.684 | 5 | 0.524 |
| 0.18 | 0.840 | 5 | 0.439 |
| 0.25 | 0.067 | 5 | 0.949 |
| 0.3548 | 0.889 | 5 | 0.415 |
| 0.5012 | 0.017 | 5 | 0.987 |
| 0.7079 | 0.202 | 5 | 0.848 |
| 1 | 1.135 | 5 | 0.308 |

### Experiment 1: Discussion

In Experiment 1, we replicated Motoyoshi and Hayakawa’s basic adaptation-induced blindness effect, ensuring that we tested equal numbers of trials at all contrast levels. It is clear from the results that this effect does not resemble classical contrast adaptation, in that the effect is equal across the entire range of target contrasts. Thus we went on to test whether it differed in other ways from classical contrast adaptation; would it transfer between the eyes, or to target stimuli at different orientations from the adapting stimulus? If, as Motoyoshi and Hayakawa suggest, the effect is a higher-level, parietal effect, it should be occurring at levels beyond binocular combination, and so we should expect to see full interocular transfer and, potentially, little or no orientation tuning.

## Experiment 2: Interocular transfer of AIB

### Methods

*Participants* Five adults with normal or corrected-to-normal vision participated in the experiment. Three participants, including two of the authors, were experienced psychophysical observers, whilst the remaining participants were naive to the purposes of the study.

*Apparatus and stimuli* Stimuli were programmed in MATLAB version 7.9 using the Psychophysics Toolbox Version 3 (Brainard 1997; Pelli 1997) Visual stimuli were presented on a Sony Triniton CPD-G500 22-inch CRT monitor with a screen resolution of 1024 × 768 pixels and a vertical refresh rate of 100 Hz. The monitor was gamma-corrected to ensure linear luminance output and was controlled by a quad core Mac Pro computer connected to a Cambridge Research Systems Bits++ digital-to-analogue converter that provided 14-bit resolution for measurement of low contrast thresholds. Maximum and minimum luminances were 67.3 and 0.26 *cd*/*m*^2^, and mean luminance was 33.8 *cd/m^2^.* Participants sat in a darkened room with their head supported by a chin rest at a distance of 57 cm from the monitor and made responses on a standard keyboard. Adapting and target stimuli were as described above, except that there were now separate displays for each eye (see Figure 2). Stimuli for this Experiment were viewed through a mirror stereoscope, with adaptors and tests presented either to the same or to different eyes.

**Figure 2.**
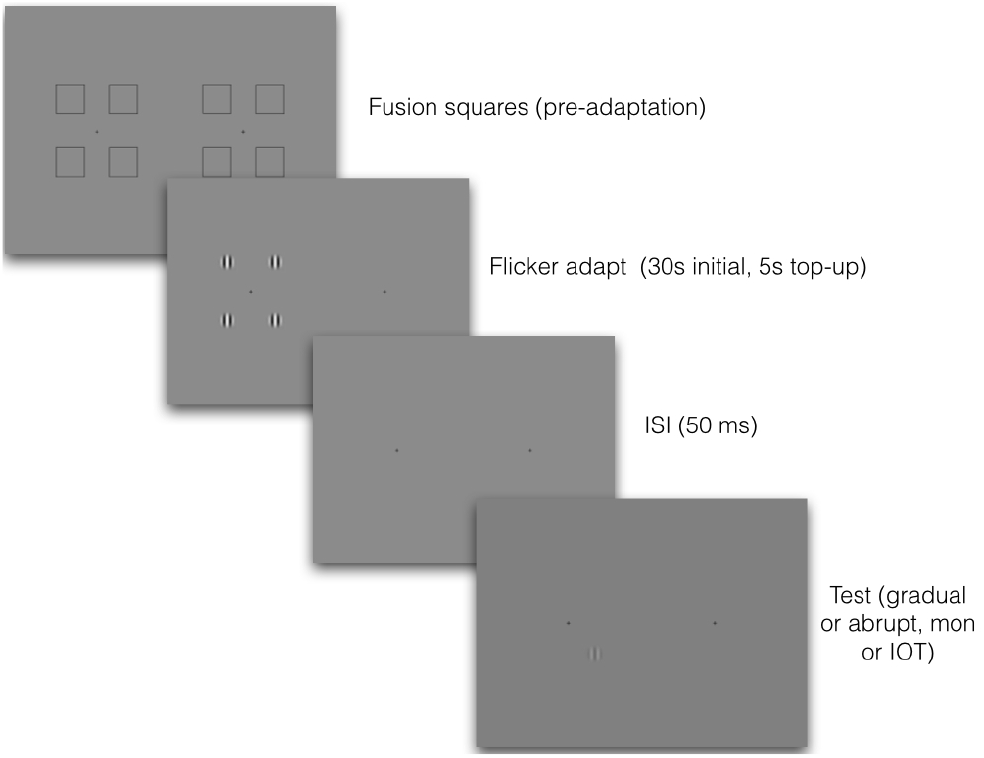
A schematic illustration of the procedure for Experiments 2 and 3. Observers first adjusted the stereoscope using the fixation squares, which then disappeared. They adapted to four counterphasing high-contrast gratings in one eye and were then tested either in the same or in the other eye, with gradual-onset or abrupt-onset target gratings. Because images were viewed through a mirror stereoscope, images on the left were seen by the left eye, and images on the right by the right eye. Target gratings were always at full contrast, and the number of incorrect judgments was recorded. Responses were always 4AFC, with participants choosing which of four locations contained the target grating.

Each adaptor appeared in a circular aperture whose diameter subtended 2° of visual angle and was edge-blurred by a cosine ramp that transitioned from minimum to maximum over 0.78° of visual angle.

The target stimulus was a static Gabor with equal spatial frequency to the adaptors but had variable orientation relative to the adaptors (0 or 90°) and temporal onset. Target stimuli were equal in diameter to the adaptors at 2^°^ of visual angle. As in Experiment 1, gradually onset target stimuli were ramped on by a temporal Gaussian with a standard deviation of 200 ms and peak amplitude at 1000 ms, while abruptly onset targets were presented within a rectangular temporal window of 300 ms duration.

To ensure that participants had sufficient stereoacuity to support interocular transfer (Mitchell and Ware 1974; Movshon et al. 1972), an Optec 2500 Vision Tester capable of testing down to 20 seconds of arc was used to test participants’ depth perception. Published thresholds for normal stereoacuity range from 40 to 60 seconds of arc (Adams et al. 2008; Romano et al. 1975). A mid-range cutoff of 50 seconds of arc was chosen because substantial interocular transfer is still observed for this level of ability (Mitchell and Ware 1974). All participants had better stereoacuity than the cut-off (M = 29, SD = 12.8).

To maximise the potential for interocular transfer, participants’ ocular dominance was determined behaviourally by a standard finger pointing measure (Coren and Kaplan 1973). Each participant pointed to a coin glued to a wall and observed how far their finger appeared to deviate when viewing with their left eye closed compared to when their right eye was closed. The open eye that caused the least deviation was judged to be dominant. Using this method, three participants were identified as right-eye dominant and two as left-eye dominant. The adaptor was always presented to the dominant eye. Several studies have shown that this method of testing produces significantly greater magnitudes of interocular transfer compared to when the non-dominant eye is adapted (Howarth et al. 2009; Mitchell and Ware 1974; Movshon and Blakemore 1973; Mohn and Van Hof Van Duin 1983).

Participants adapted to counterphasing stimuli presented to either the left or right eye. In monocular conditions, the adapting stimulus was presented to the non-dominant eye. In order to maximise the potential for interocular transfer to occur, participants had their dominant eye adapted in the interocular conditions. This meant that the target was always presented to the non-dominant eye, regardless of ocular condition.

Participants pressed a key to initiate trials and were presented with four counterphasing adaptor gratings, one appearing in each aperture. Adaptors were displayed for 30 s for the initial two trials, with 5 s ‘top-ups’ for subsequent trials. (Pilot testing with 60s initial adaptation and 10s top-ups indicated that the shorter adaptation times produced identical results). Following offset of the adaptors, a spatial four-alternative forced-choice task required observers to use a standard keyboard to indicate which aperture the target appeared in. Conditions were blocked by target orientation, temporal onset of the target (gradual versus abrupt), and tested eye (monocular vs. IOT conditions). The order of completion was counterbalanced to ameliorate practice effects.

### Results: Experiment 2: Interocular transfer and orientation specificity of AIB

In this experiment, we presented all the target stimuli in the 4AFC paradigm at full contrast, either ramped up gradually or with abrupt onset, and measured the proportion of times the observers failed to correctly identify the location of the target grating. Participants’ subjective reports indicated that in many instances, the gradual onset targets were simply not seen, even though they were ramped up to full contrast. This is shown clearly in the data (see Figure 3); target disappearances are strongly orientation-specific, but interestingly, there was absolutely no interocular transfer of the effect. In addition, abruptly presented targets never disappeared (perhaps unsurprisingly, since they were at full contrast).

**Figure 3.**
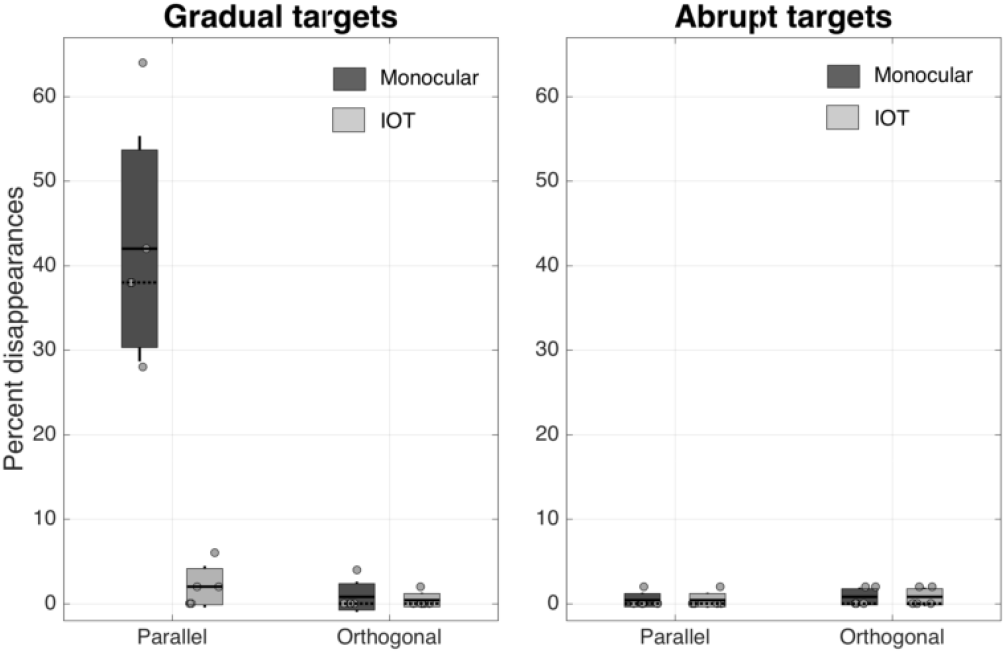
Percentage of disappearances of full contrast target stimuli after adaptation to counterphasing gratings, for target stimuli which were either 0 degrees or 90 degrees of relative orientation from the adapting stimuli, with either gradually onset (a) or abrupt (b) target stimuli, tested either in the same eye as the adapting stimulus (dark grey) or in the other eye (light grey). Shaded areas 1 standard error, lines show standard deviations, and individual grey dots are individual data points. Solid lines show the means, and dotted lines the medians. All five subjects completed all conditions.

There were significant main effects of onset type (gradual vs. abrupt), *F*(1, 4) = 58.82, p = .002, eye tested (monocular vs. IOT), *F*(1,4) = 35.36, p = .002, and relative orientation (0 vs. 90 degrees), *F*(1,4) = 50.69, p = .002. There were also significant two-way interactions between onset type and eye tested, *F*(1,4) = 35.36, p = .004, onset type and relative orientation, *F*(1,4) = 68.48, p = .001, and eye tested and relative orientation, *F*(1,4) = 32.83, p = .005. In addition, there was a significant three-way interaction between onset type, eye tested and relative orientation, *F*(1,4) = 32.83, p = .005. In summary, the only case in which adapting to the flickering pattern caused disappearance of the target was in the monocular condition where the gradually presented target was parallel to the adaptor, as can be clearly seen from Figure 3.

## Experiment 3: Orientation tuning of AIB

We further measured the orientation tuning of the AIB disappearances in more detail (see Figure 4), since orientation tuning of this effect was not reported in the original paper. Here it is clear that the effect is tightly orientation tuned, and is well fitted by a Gaussian function with a standard deviation of 7.76° and an amplitude of 37.4%. The fitted amplitude of the function (37.4%) is very close to the approximate percentage of target disappearances at 0 degrees, on average, across participants. We note that this is also very close to the reported percentage of incorrect responses (34%) in the original AIB paper (Motoyoshi and Hayakawa 2010). It should be noted that the actual percentage of targets not seen in our experiment (and in Motoyoshi’s, in fact) would probably be higher, given that the 4AFC paradigm would result in a 25% guess rate, on average. It is important to note here that instances of disappearance were almost completely absent at full contrast, regardless of the relative orientation of adaptor and target. The implications of this will be covered in the Discussion.

**Figure 4.**
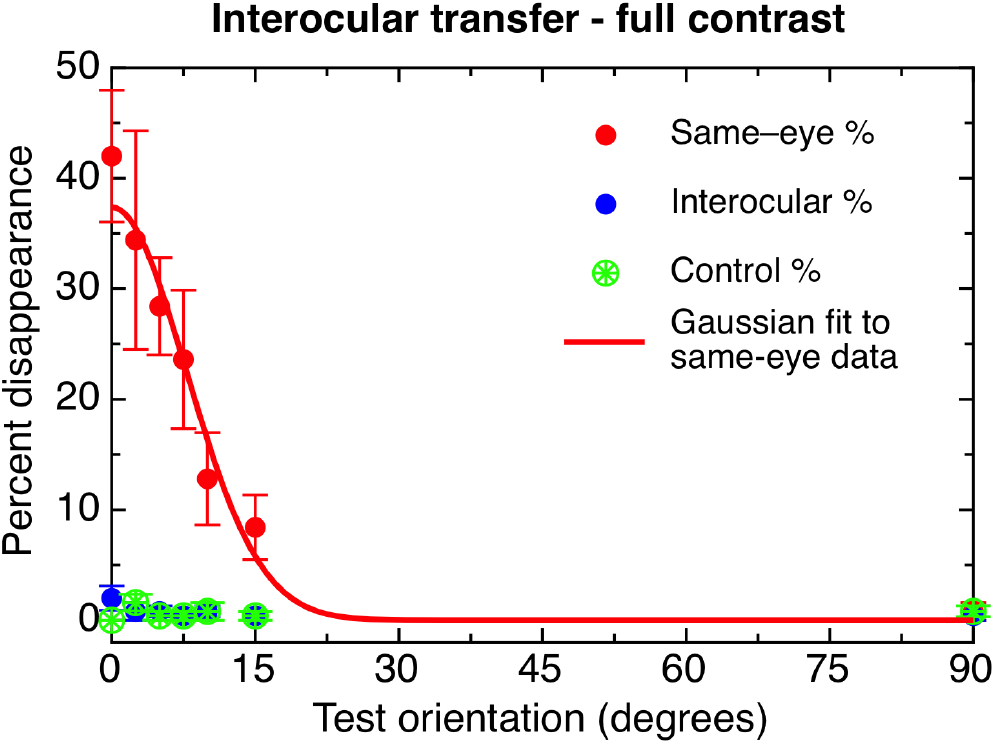
The orientation tuning of disappearances (measured by error rates) at full contrast. Red data points show percent disappearances for stimuli presented in the same eye as the adapting stimuli; blue points show those presented in the unadapted eye, and green show the control stimuli presented without adaptors. The solid line shows a Gaussian fit to the data, fitted in ProFit Version 6.2.11, using a Leverburg-Marquardt algorithm; the fit was given two free parameters (amplitude and bandwidth), with a fixed mean of 0 and baseline of 0. Standard deviation of the fit was 7.76 degrees, with an amplitude of 37.4%.

## Discussion

The first demonstration of AIB at the 2008 Vision Sciences Society met with astonishment. The audience was shown two high contrast Gabor patches: one with an abrupt temporal onset profile, the other a more gradual profile. Unsurprisingly, both were trivially easy to detect. A flickering Gabor stimulus followed, and the audience was instructed to fixate upon it for about ten seconds. This adapting stimulus was then replaced with the high contrast targets. To the audience’s audible surprise only one patch was visible – the abrupt one. The gradual patch was completely invisible, apparently suppressed from awareness. Why had it disappeared? Was it a form of threshold elevation? What neural mechanisms might be responsible? To help understand this phenomenon, we replicated this basic effect across a range of very specific conditions. Experiment 1, showed that AIB effects are indeed profound, with gradual target detection performance plummeting to chance levels at all contrasts tested (0.06–1.0 contrast).

Experiments 2 and 3 were designed to measure AIB’s selectivity for both orientation and interocular transfer. Our results show very clearly that the AIB effect is tightly selective for orientation. Disappearances were common when the adaptor and target Gabors were similarly oriented, with performance improving monotonically out to approximately 20 degrees of relative orientation, at which point performance reached ceiling (100% accuracy). That AIB should prove so specific in the orientation domain implicates that it is mediated by a population of neurons with a similarly tightly tuned orientation preference. Neurons with such receptive fields are common in several early visual cortical areas, including areas V1, V2 and V3 (Hubel and Wiesel 1968; Boynton and Finney 2003). Exactly how early in cortical processing are the neurons which mediate AIB? Experiments 2 and 3 addressed this with an interocular transfer manipulation. While disappearances were common when adapting and testing in the same eye, adapting a single eye to flicker and presenting the target to the other eye afforded nearly perfect performance. Such a high degree of eye specificity shows that AIB is purely monocular.

From a classical view of adaptation, our finding that AIB is both orientation-tuned and purely monocular is surprising. It is now well established that threshold elevation, which occurs as a consequence of flicker adaptation, is composed of a binocular and a monocular component (Baker and Meese 2012; Cass et al. 2012). The binocular component is orientation-tuned whilst the purely monocular component is untuned. That AIB should be both tightly orientation tuned and purely monocular distinguishes it, therefore, from classical adaptation-induced threshold elevation. It also suggests that AIB involves a subset of early cortical neurons whose receptive fields are both orientation specific and purely monocular. fMRI evidence indicates that both orientation-specific adaptation and eye of origin information and is evident in early visual cortical areas, V1, V2 and V3 (Boynton and Finney 2003; Schwarzkopf et al. 2010) although purely monocular neurons tend to be found only in V1.

One study investigating the precise relationship between neural orientation and eye preference in primate V1 (Bartfeld and Grinvald 1992) found that orientation-selective cells are in fact located within ocular dominance columns. Given that ocular dominance columns contain neurons which are both orientation tuned and ocularly specific in their response, it seems reasonable to speculate that AIB may therefore be mediated by these cells, located close to the centre of a single ocular dominance column, such as those found in V1 (Bartfeld and Grinvald 1992).

This conjecture that AIB occurs relatively early in the visual cortical process is supported by two visual crowding studies showing that crowding can be largely extinguished if the flanking stimuli which produce the crowding effect are suppressed from awareness using AIB (Wallis and Bex 2011; Shin and Tjan in press). According to Shin and Tjan (in press), this implies that AIB is likely to occur at a level of neural processing preceding visual crowding. fMRI studies indicate that crowding effects can be differentiated as early as V1 (Anderson et al. 2012). If AIB does in fact precede crowding, this would point to AIB being mediated at a very early stage of visual processing.

Motoyoshi and Hayakawa’s 2010 AIB effect is undoubtedly a striking and curious phenomenon that remains to be fully understood. In first reporting the phenomenon, the authors? original conjecture was that AIB occurred relatively late in visual processing. The evidence presented here, however, from our orientation and interocular transfer experiments and complemented by related work on AIB and crowding, converge on the conclusion that AIB is very likely to be mediated at a very early cortical stage of visual processing.

## Acknowledgements

David Alais would like to acknowledge ARC Discovery Project DP0878371

John Cass would like to acknowledge ARC Discovery Project DP0774697

